# Food deprivation modulates heart rate, motor neuron, and locomotion responses to acute administration of d-amphetamine in zebrafish larvae

**DOI:** 10.1101/2022.05.31.494235

**Authors:** Pushkar Bansal, Mitchell F. Roitman, Erica E. Jung

## Abstract

Psychostimulant drugs are so named because they alter the cardiac, brain and behavioral responses in humans and other animals. Acute food deprivation or chronic food restriction potentiates the stimulatory effects of abused drugs and increases the propensity for relapse to drug seeking in drug-experienced animals. The mechanisms by which hunger affects cardiac and behavioral activities are only beginning to be elucidated. Moreover, changes in motor neuron activities at the single neuron level induced by the stimulants, and their modulation by hunger, remain unknown. Here we investigated how the state of hunger affects responses to d-amphetamine by measuring locomotion, cardiac output, and individual motor neuron activity in zebrafish larvae. We used wild-type larval zebrafish to record behavioral and cardiac responses and the larvae of mnx1:GCaMP transgenic zebrafish to record motor neuron responses. Acute administration of d-amphetamine in sated larvae did not induce a significant change in the motor responses (swimming distances, tail activity), heart rate, or motor neuron firing frequency to the stimulant. However, food deprivation enhanced amphetamine-evoked responses significantly. The results extend the finding that signals arising from food deprivation are a key potentiator of the drug responses induced by d-amphetamine to the zebrafish model. The larval zebrafish is an ideal model to further elucidate this interaction and identify key neuronal substrates that may increase vulnerability to drug reinforcement, drug-seeking and relapse.

## Introduction

Drug abuse in general and amphetamine abuse, in particular, remain pressing medical and societal concerns. A greater understanding of factors contributing to the psychomotor activating effects and the ability of drugs to reinforce behavior is critical for the development of effective interventions. One factor that influences amphetamine drug reactivity, self-administration and relapse is food restriction. Acute food deprivation or chronic food restriction enhances the locomotor stimulating effects of amphetamine.^1,2^ Food restriction increases the rate of acquisition for drug self-administration as well as drug-seeking in abstinence and reinstatement paradigms.^3,4^ In humans, a twenty-four hour fast enhances amphetamine discriminability.^5^ Taken together, these data support food restriction as a risk factor for the development of drug seeking and continued drug seeking in individuals with a history of drug taking. While much debate remains, drug and food rewards compete with one another in food restricted animals.^6^ Thus, understanding the interaction between food restriction and psychomotor interaction is of critical importance.

Zebrafish (*Danio rerio*) larva is a widely used animal model in neuroscience research. Being optically transparent, zebrafish larva makes an exceptional research model for imaging and controlling neuronal activity and measuring other internal processes such as heart rate and blood flow in response to external stimuli via the optical window. Zebrafish larva is a newly emerging animal model used in abused drug studies and has been adapted in a limited number of studies to observe locomotion in response to stimulant drugs. Acute administration of amphetamine and ethanol leads to hyper-locomotor activity in zebrafish larvae.^7^ Locomotor response to antipsychotic drugs has also been evaluated by researchers in zebrafish larva.^8^ However, to the best of our knowledge, there are no studies investigating the modulation of drug responses in zebrafish larvae under sated and food restricted states. Responses of zebrafish larvae to psychostimulants under fed and fasted states would open up a new possibility of using zebrafish larvae as a comprehensive animal model to study the distinct neural substrates and processes whereby circuitry critical for homeostasis interacts with motor circuits critical for abused drug responses. Results could pinpoint where access and control of individual neuronal activity can serve as an intervention.

In this paper, we aim to study whether hunger affects drug responses to acutely administered stimulant (d-amphetamine; AMPH) in zebrafish larvae. We delivered AMPH to zebrafish larvae in one of two physiological states: sated and fasted. Motor responses (e.g. locomotion and tail activity) and heart rate were continually monitored. In addition, individual motor neuron activity was recorded in real-time before and after AMPH administration. Our analysis indicates that zebrafish larvae in the fasted state exhibit a significant increase in behavioral and physiological responses relative to those in sated states post AMPH administration. Overall, our findings point to the difference in behavioral, physiological, and neuronal activities in response to AMPH given under sated and starved states.

## Results

### Experimental groups and performed analysis

In this study, we used 6-7 days post fertilization (dpf) zebrafish larvae to observe the effect of the stimulant (d-amphetamine hemisulfate (AMPH)) on motor activities such as total distance travelled, heart rate, and motor neuronal activity. Distance travelled by larvae was recorded in a 48-well plate, whereas heart rate and neuronal activities were recorded by placing larvae in a petri dish. Details on experimental settings and imaging conditions are in the methods section. We divided zebrafish larvae into four experimental groups, fed control (fed larvae without AMPH), food-restricted control (food-restricted larvae without AMPH), fed + AMPH (fed larvae treated with AMPH) and food-restricted + AMPH (food-restricted larvae treated with AMPH) (Fig 1A). Food-restricted groups were kept food-deprived from birth, and fed groups were given food daily once they turned 5 dpf following a normal feeding schedule. Food-restricted and fed groups of larvae were kept in separate petri dishes until the experiment. The former two groups (fed control and food-restricted without AMPH exposure (- AMPH)) together constitute control groups, and the latter two groups (fed and food-restricted with AMPH exposure (+ AMPH)) together account for AMPH treated groups.

**Figure 1.**
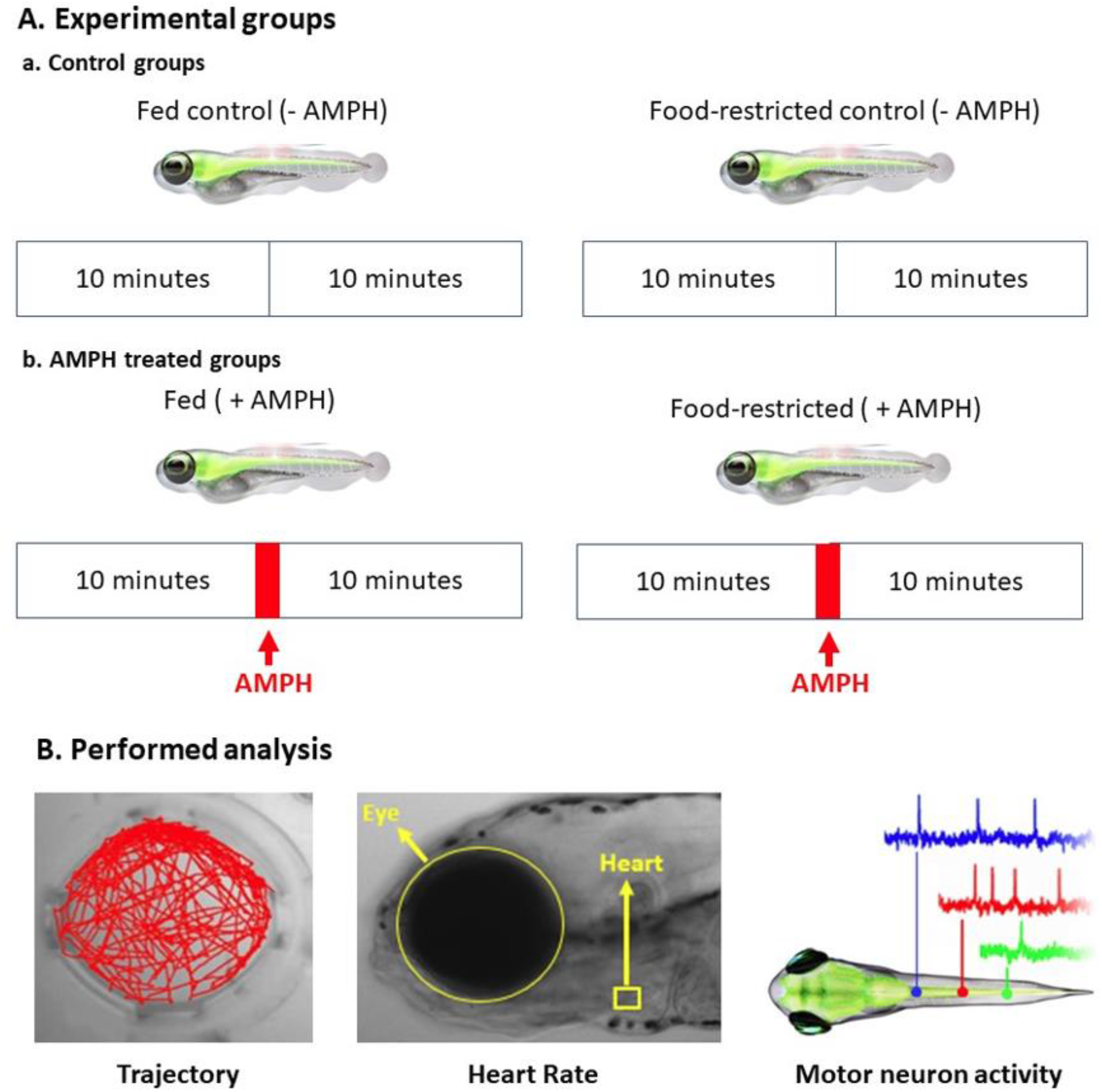
Experimental groups and Activities. A) Four experimental zebrafish larval groups were used under fed and food-restricted states to observe the behavioral, physiological, and neuronal responses to stimulant-amphetamine (AMPH). Fed control and food-restricted control were control groups that were recorded without AMPH. Fed with stimulant and food-restricted with stimulant were AMPH treated groups that were stimulated using AMPH only after 10 mins of recording without stimulant. Recording continued shortly after for 10mins after the AMPH was administered. B) Responses in zebrafish larvae to AMPH were analyzed with its three different attributes, locomotion, heart activity and motor neuron activity.

Control groups of zebrafish larvae (fed and food-restricted) were recorded for 20 mins without any AMPH administration. Activities in AMPH treated groups (fed + AMPH and food-restricted + AMPH) were recorded for the first 10 mins in 0.5ml of fish water in the wells without AMPH, followed by pipetting out the fish water and adding 0.5 ml of 0.7uM d-amphetamine into the wells. We waited for 2-3 mins to allow the larvae to absorb the AMPH through its skin and mouth, and the recording was continued again for 10 more minutes to observe motor activities (distance travelled) and heart rate. For motor neuronal activity, recordings were conducted for 20 minutes before and after the AMPH treatment. Figure 1B shows the four quantitative assays performed in this study, distance travelled by zebrafish larvae, heart rate activity and motor neuron activity in the larval spinal cord, to understand their changes in response to AMPH under fed and food-restricted states.

### Hunger potentiates locomotor activity in the presence of d-amphetamine

To observe changes in locomotor activity in freely swimming zebrafish larvae, we used a customized apparatus to visualize and measure the locomotor response (total distance travelled), as shown in Fig.2A. Details of the imaging setup and image acquisition are in the method section. We measured the total distance moved by zebrafish larvae in control and AMPH-treated groups for 20 minutes (in 10 minutes segment as shown in Fig. 1A). Figure 2B shows representative trajectories of larval motion observed for 10 minutes before and after the AMPH administration. Visual observation of larval motion trajectories in the wells before and after AMPH exposure showed hyperactivity after AMPH treatment in both food-restricted and sated larval groups (Fig. 2B).

**Figure 2.**
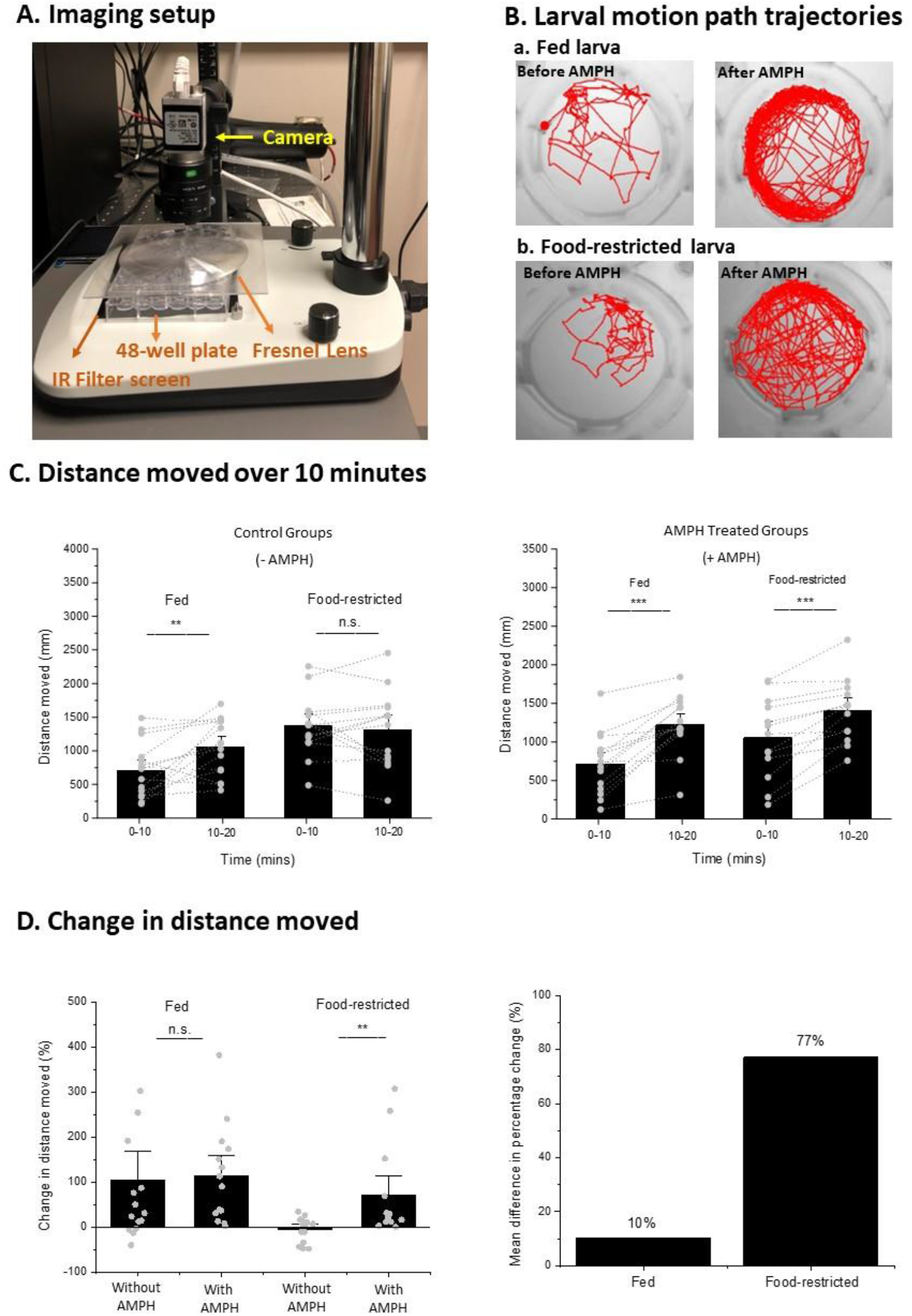
Motor activity imaging setup and locomotor response. A) Customized setup for larval motor activity imaging and recording that consisted of a high-speed camera mounted on a camera stand. A dissection stereoscope with removed eyepiece and its light source was used to illuminate the 48-well plate through an IR filer screen, and a Fresnel lens screen was placed over the well plate to minimize image distortion. This setup was utilized to record locomotor activity. B) The imaging setup described in Fig. 2A was connected to Noldus Ethovision XT 15, an activity tracking software that created the motion trajectories of zebrafish larvae before and after AMPH administration in fed and food-restricted groups. C) Locomotor activity was recorded for a total of 20 min time period (10-min binning). Fed control group (n=15) showed heightened locomotor activity after 10 mins with no AMPH administered, and food-restricted control group (n=14) activity remained unchanged (paired t-test (normal distribution), p<0.01). d-amphetamine caused the locomotor activity to increase in both fed (n=14) and food-restricted (n=13) states, with overall activity remaining relatively higher in the food-restricted state (paired t-test (normal distribution), p<0.0001). D) Mean difference in percentage change was calculated between the first and 10 mins and the next 10 mins of locomotor activity shown in Fig. 2C for control and AMPH treated groups. These percentage change differences when compared between control and AMPH treated groups in their respective fed and food-restricted states, showed an increment of 10% and a significant rise difference in the food-restricted state by 77% (Mann-Whitney test (skewed distribution), p<0.01). Statistical comparison bars are represented as standard error of the mean (SEM) bars.

Figure 2C shows distances travelled during the 10 minutes segment by the control (- AMPH) group and AMPH treated (+ AMPH) group. In control groups without AMPH exposure, fed (n=15) larvae displayed a significant increase (p < 0.01) in total distances moved, while food-restricted larvae (n=14) showed an insignificant change in locomotor activity when comparing the first 10 minutes and the next 10 minutes. We repeated recordings of moving distances in AMPH treated groups, this time with brief AMPH exposure between the first 10 mins and the next 10 mins of recordings. In both fed (n=14) and food-restricted (n=13) groups, locomotor activity was increased significantly (p < 0.0001) after the AMPH was administered. To understand the effect of the AMPH on locomotor activity in fed and food-restricted groups, we calculated the percentage change in locomotor activity in fed and food-restricted larvae by comparing the distance moved during the first and next 10 minutes before and after the AMPH administration (Fig. 2D). We observed an increase in the locomotor activity of fed larvae in both control (without AMPH) and AMPH treated (with AMPH) groups (n = 15 and n=14, respectively) when comparing the first and following 10 minutes of locomotion (Fig. 2D). The relative change between with and without AMPH treatment groups under the fed state was found to be insignificant (10%) (Fig. 2D). However, in food-restricted larvae, the control group (without AMPH) exhibited an insignificant decrease in activity, whereas the AMPH treated group (with AMPH) exhibited a significant increase when comparing distances moved pre-and post-AMPH treatment (n = 14 and n=13, respectively). Figure 2D shows the mean difference between percentage change that was calculated for fed and food-restricted groups without and with AMPH over 10 min interval. Mean difference was significantly high in food-restricted group with an increment of 77% (p < 0.01) whereas the percentage difference for fed group only increased by 10% after AMPH exposure. Overall, the change in locomotor activity was higher after AMPH exposure in food-restricted group when compared to the change occurred in locomotion in fed group.

AMPH exposure can affect other motor behavior-related factors such as higher/lower time periods of activity and moving speed (or impulse). To assess these factors, further behavioral evaluation was performed, where we compared the total amount of active moving period of larvae (i.e., when larvae made tail movement at least once in 10 second time period during the continuous recording). Moving speeds (i.e., total moved distance divided by the active period) were compared in fed and food-restricted groups with and without AMPH treatment in fed and food-restricted states. A previous study demonstrated that d-amphetamine lowered impulsive behavior in sated rodents.^9^ As a representative of impulsive behavior in zebrafish, we assessed the swimming velocity change over time. Figure 3A shows representative pictures of distance moved per 10s during the 10 minutes period before and after AMPH exposure, where a sated larva moved less impulsively than a food-restricted larva after the AMPH exposure (Fig. 3A).

**Figure 3.**
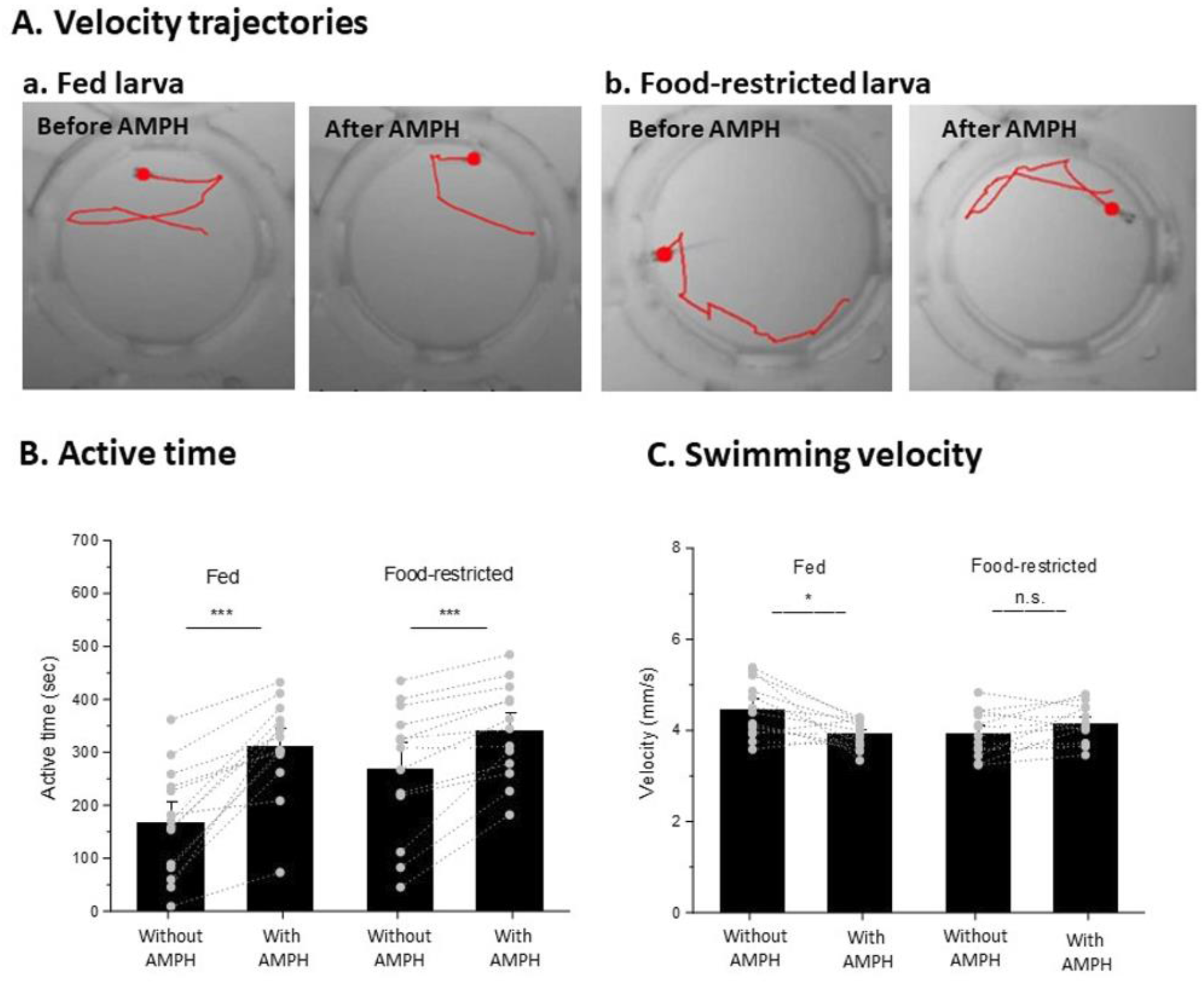
Active time and impulsive behavior from velocity change. A) Pictorial representation of fed and food-restricted larva portrays the distance travelled by the larva in a 10-sec time period before and after AMPH administration as velocity. From these velocity trajectory illustrations, the fed larva was observed to move with lower velocity, which can also be interpreted as a movement with lesser impulsivity exhibited by the fed larva after the AMPH was given. B) Both fed and food-restricted larvae were found to be significantly active for longer periods after AMPH exposure (paired t-test (normal distribution), p<0.0001). C) Statistical comparison of velocity change after AMPH administration (impulsivity) shows a marked decrease in fed larval groups whereas food-restricted group were not to be impulsive after AMPH administration (Fed group (n=14): paired Wilcoxon signed rank test (skewed distribution), p<0.01; Food-restricted group (n=13): paired t-test (normal distribution), p>0.05, n.s.)). Statistical comparison bars are represented as standard error of the mean (SEM) bars.

**Figure 4.**
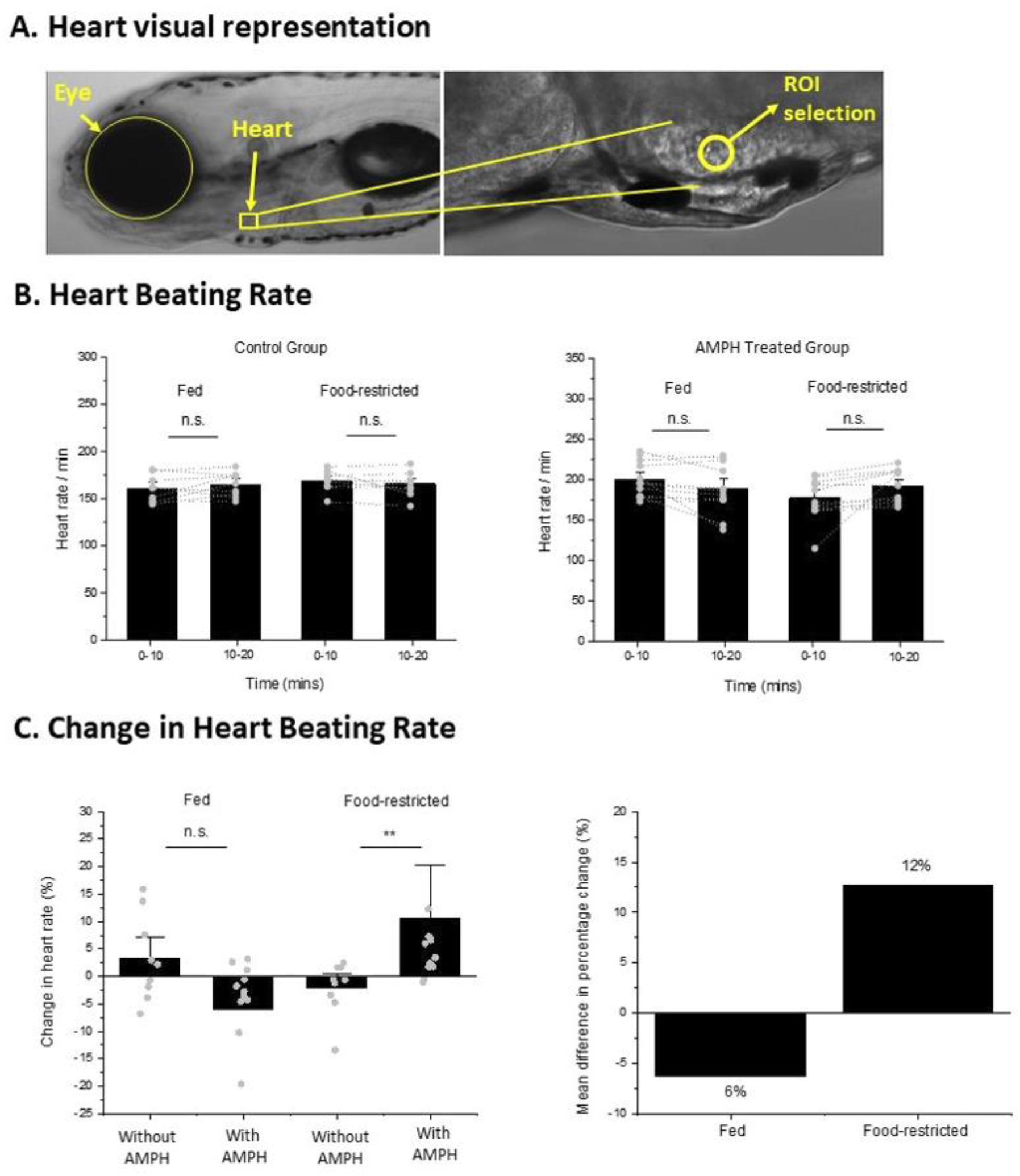
Heart rate response to d-amphetamine. A) The picture shows the exact location of the larval heart with respect to other physical parts and organs. Heart rate activity in zebrafish larva was recorded with a high-speed sCMOS camera mounted on an inverted bright-field microscope and recorded with HCI image live software. Region of interest (ROI) was selected (avoiding noisy regions for accuracy), and heart activity measurements were performed using ImageJ. B) Heart rate activity was not found to be significant in the absence and presence of d-amphetamine under fed (n=9 for control, n=12 for AMPH treated) and food-restricted states (n=9 for control, n=12 for AMPH treated) (paired t-test (normal distribution), p>0.05, n.s.) although, minor decrement in the fed group and increment in the food-restricted group can be observed in heart rate after d-amphetamine was administered. C) Percentage change difference comparison shows a difference decrease (6%) in heart rate in fed groups of control and AMPH treated larvae, but a significant difference increase (12%) in heart rate was found in food-restricted groups (Mann-Whitney test (skewed distribution), p<0.01). Statistical comparison bars are represented as standard error of the mean (SEM) bars.

To calculate swimming velocity, active time periods were isolated by excluding time periods when larvae were immobile during the 20 minutes of recording. Figure 3B shows that the active swimming period increased in both fed and food-restricted groups after AMPH exposure (n = 14 and 13, respectively, p < 0.0001). Then swimming velocity was calculated by dividing the total moved distance by the active swimming time. Figure 3C presents that fed larvae showed a 15% decrement (p<0.01) in velocity when exposed to the AMPH, whereas food-restricted larvae exhibited a 6% increase in velocity in response to the AMPH (Fig. 3C). Data shows that fed larvae moved with a decreased velocity after being exposed to AMPH than before, whereas food-restricted larvae moved with increased velocity. Overall, our findings in locomotor behavior in free-swimming larvae indicate that AMPH exposure intensified locomotion in the larvae in the food-restricted state that moved greater distance with an increased swimming velocity compared to the larvae in the sated state that moved shorter distances with a decreased swimming velocity.

### Heart rate is affected contrastingly in d-amphetamine’s presence

According to a clinical study in humans, cardiac response changes with psychostimulants administration in the food-restricted and sated state (Angrist, 1987), and we investigated how it gets altered in zebrafish larvae under both physiological states. To measure the changes in heart rate and motor neuronal activity in zebrafish larvae, we used an inverted bright-field microscope equipped with a 40x water immersion objective and a high-speed sCMOS camera to record heart beating rate. Heart rate requires the larvae to be immobilized; thus, larvae were fully immobilized in a 1.5% agarose gel drop. Details of the experiment are elaborated in the method section. Figure 5A presents a visual description to show the location of the heart in zebrafish larvae and the ROI, which we selected to measure heart beating rate using opensource software (ImageJ). We carried out an initial evaluation of heart rate for the four groups (fed and food-restricted larvae with and without AMPH treatment) and did not find any significant difference in fed and food-restricted larvae in control (n=9 for fed and food-restricted larvae) and AMPH treated groups (n=12 for fed and food-restricted larvae) (Fig. 5B). When comparing changes in heart beating rate between the first and next 10 mins, fed larvae showed a decrease in heart beating rate after AMPH treatment (Fig. 5C). Conversely, food-restricted larvae displayed an incremental change in heart beating rate after AMPH treatment (p<0.01). Mean differences in percentage change between fed and food-restricted states without and with effect of AMPH exhibited a decline in by 6% in the fed state, while food-restricted larvae exhibited an increase in percentage change difference in response to AMPH by 12%. This result was found be to similar to locomotor activity as the change in heart rate post-AMPH administration surged in food-restricted larvae contrary to the change in fed larval group where the percentage change exhibited a decreasing trend.

**Figure 5.**
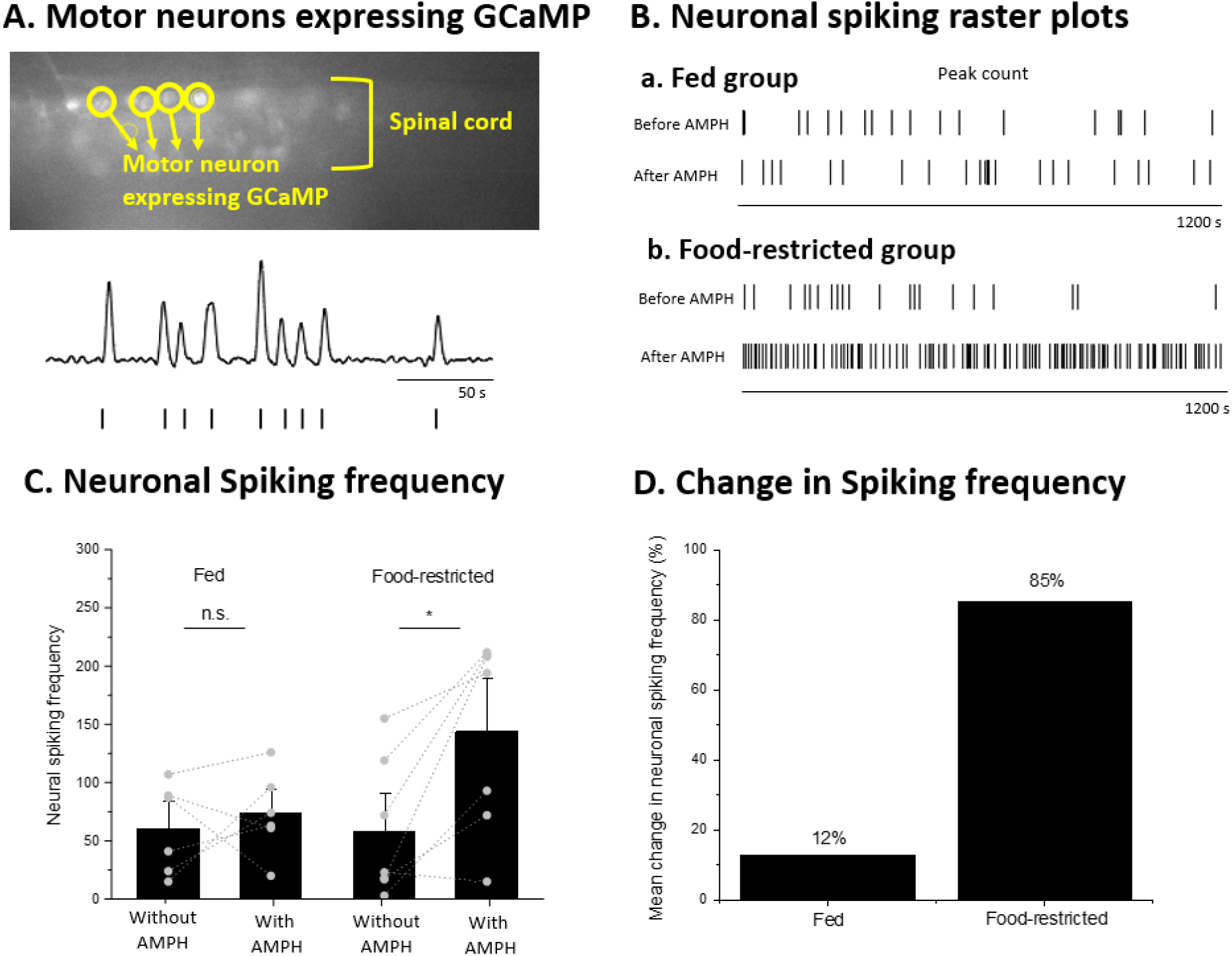
Motor neuronal activity change response to d-amphetamine. A) Activity in motor neurons present in the spinal cord of the larval spinal cord affects locomotor activity. The pictorial representation of paralyzed and completely immobilized zebrafish larva (mnx1:GCaMP strain) shows the motor neurons that express GCaMP, a calcium indicator that emits fluorescence after excitation. Neuronal activity was recorded using an epifluorescent microscope connected to a high-speed sCMOS camera and recorded using HCI image live software. Motor neuronal ROIs selection and activity measurement was performed using ImageJ. B) Raster plots generated from neuronal activity shows neuronal spiking count in fed and food-restricted groups before and after AMPH administration over 1200 sec (20 mins) time period. These raster plots are a visual representation that shows the magnitude of change in neuronal activity post AMPH administration. C) Neuronal spiking frequency/count increased in both fed (n=6) and food-restricted (n=7) groups after larvae were exposed to d-amphetamine, but significance can only be observed in the food-restricted state (paired t-test (normal distribution), p<0.05). D) Percentage change in neuronal activity induced by d-amphetamine was found to be 12% in the fed group and 85% (significantly high) in the food-restricted state. Statistical comparison bars are represented as standard error of the mean (SEM) bars.

### Activity in motor neurons is affected by d-amphetamine in hungry state

Locomotion is controlled by motor neuronal activation, which causes muscle movements by generating action potentials that send locomotory signals to muscle fibers. We hypothesized amphetamine to invoke hyperactivity in motor neurons, which results in hyperactivity in larval motion. To measure neural activity response in individual motor neurons *in vivo* zebrafish larvae, a transgenic line, *Tg* (mnx1:GCamP), that expresses a genetically encoded calcium indicator (GCaMP) in motor neurons was used in this study.^10^ To avoid unnecessary motion of the larvae during recording, larvae were paralyzed with 20ul of 300uM pancuronium bromide (nondepolarizing muscle relaxant (Hong, 2016). Fluorescence images of motor neurons expressing GCaMP were imaged under an epifluorescence microscope equipped with an sCMOS high-speed camera (Fig. 6A). Details of paralysis protocol and imaging conditions are in the methods section. Motor neuronal activity in zebrafish larvae was recorded at 35 frames per second for 20 minutes pre- and pose AMPH exposure. Figure 6B shows raster spike plots to represent neuronal spiking frequency counts before and after AMPH administration in food-restricted and fed larvae over 20 minutes. In a fed group, no noticeable difference was observed between spiking counts before and after AMPH administration. However, we observed significantly increased spiking counts after AMPH administration in food-restricted groups. When we compared the neuronal spiking frequency over 20 minutes (Figure 6C; n=6 and n=7 for the fed and food-restricted states, respectively), no significant change was observed in larvae under fed conditions. However, neuronal activity increased significantly (p < 0.05) after AMPH treatment in the food-restricted larval group. From a differential percentage change analysis, on an average, food-restricted larvae exhibited an 85% increase in spiking frequency after AMPH exposure, while fed larvae showed a 12% increase (Fig. 6D).

## Discussion

To investigate the effect of hunger and satiety on drug reactivity, we have studied behavioral, physical, and neuronal responses to the psychostimulant drug (d-amphetamine) in zebrafish larvae. Abused stimulant drugs have been shown to adversely affect behavioral and brain activities in humans and animals.^11,12^ On the other hand, physiological states such as hunger and satiety are well known to substantially impact behavioral and brain activities.^13–15^ However, the gap of knowledge remains unplugged in the understanding of how behavioral and neuronal responses to abused stimulant drugs are altered under hunger and satiety states. In this pre-clinical study, we tried to fill this gap by evaluating various drug responses (locomotion, heart beating rate, and neural activity) to understand the combined effects of different physiological states and stimulant drug on these responses in zebrafish larvae.

To carry out this study, we administered d-amphetamine hemisulfate to 6-7dpf zebrafish larvae in the fed and food-restricted states and compared how their responses get altered when exposed to d-amphetamine. In previous studies in rodents, the impact of a food-restricted state and exposure to a stimulant drug has been studied independently, and the individual factors increased behavioral activities in rodents.^1,2,16^ These reports were found to be consistent with our findings when considering the effect of a single factor, hunger or psychostimulant drugs. When the effects of hunger and psychostimulantwere combined, we found that the food-restricted state enhanced the effect of amphetamine, which led to a significant increase in total moving distances compared to the well-fed state, where we observed an insignificant increase in total distance moved. Changes in swimming velocity, which can be interpreted as impulsive behavior, also showed a difference between fed and food-restricted state larvae. Larvae in the fed state showed a decrease in swimming velocity after the amphetamine exposure compared to the food-restricted larval group, where an increase in swimming velocity was observed. Cardiac output is another crucial response that could impact an animal’s behavior. Studies have shown that behaviors such as aggression, violence, depression and anxiety are related to changes in heart rate.^17–19^ Pre-clinical studies in rodents also provide evidence of the effect of amphetamine on heart rate.^20^ Nevertheless, our knowledge regarding cardiac activity in response to stimulant drugs under different physiological states is limited. To fill the gap, we measured changes in heart rate in response to d-amphetamine and observed, under the food-restricted state, heart rate increased in food-restricted larvae after amphetamine administration while fed larvae exhibited a decreasing trend.

Monitoring motor neuron activity is critical to tracing the underlying neural pathway by which the reward circuitry conveys information to alter motor outputs. A number of studies observed the impact of stimulant drugs on neuron activity.^21–23^ However, a limited number of studies investigated the effect of stimulant drugs on motor neuron activity due to the technical limitation (e.g., the presence of motor neurons deep inside the spinal cord hinders the reach of monitoring probes in such hard-to-reach spots, the opacity of mammalian body). Zebrafish larvae offer an ideal animal model to observe individual motor neuron activity due to their optical transparency and densely packed motor neurons along the spinal cord. We visualized individual motor neuron activity using a fluorescent neural activity indicator (GCaMP) and observed changes in the fluorescence intensity corresponding to individual neuron activity. For the first time, we observed that starvation dramatically increases the response of motor neuron activity to d-amphetamine.

There have been studies to understand how hunger and satiety affect drug response by investigating relevant hormones and neural circuits. Interactions between hunger-mediating hormones and drug reactivity are relatively well known.^24^ Previous studies demonstrated an increased level of ghrelin, the appetite-increasing hormone and enhanced behavioral response to abused drugs.^25–29^ In contrast, an increased leptin level (the appetite-suppressing hormone) demonstrated attenuated relapse to heroin seeking.^4,30^ In contrast to the hunger-mediating hormones, the interaction between hunger-mediating neurons and drug response remains unknown. Multiple neuronal populations, such as AgRP/NPY neurons in the hypothalamus, are known to regulate appetite.^31,32^ A recent study demonstrated that the ablation of AgRP/NPY neurons in prenatal or early postnatal mouse brain resulted in more voracious drug-seeking behavior compared to untreated mice.^33^ A subsequent study demonstrated that the activation of AgRP/NPY neurons decreased anxiety levels, which can appear controversial to the contemporary view that hunger enhances drug response that is known to be positively related to anxiety levels.^34^ Hence, further studies involving the roles of appetite-controlling neurons in the presence of stimulant drugs are in need to understand behavior and uncover neuronal pathways via which neuronal populations are connected to the reward system.

## Conclusion

In summary, we have shown how satiation and food deprivation in 6-7dpf zebrafish larvae affects the behavioral, physiological and neuronal responses to a psychostimulant, d-amphetamine. We observed higher motor activities in food-restricted larvae than those in the satiated state in response to the stimulant drug exposure. In addition to this, we observed less impulsive behavior in sated larvae in response to amphetamine than those in food-restricted larvae. Heart rate in sated larvae tends to drift down over time while increasing over time in food-restricted larvae, which could be an attribute of amphetamine. Finally, we looked at the response of individual motor neurons to amphetamine exposure. Similar to locomotor response, zebrafish larvae under both sated and food-restricted states showed heightened neuronal activity response to d-amphetamine, but food-restricted larvae exhibited significantly higher neuronal activity than sated larvae. Hence, from this study, we conclude that hunger/starvation enhances motor activity, heart rate and activity in the motor neurons of zebrafish larvae in response to stimulant drugs.

## Experimental Methods and materials

### Zebrafish maintenance and breeding

Adult zebrafish were maintained in the zebrafish facility housed at The University of Illinois at Chicago in a fish racking system (Aquaneering, Inc., San Diego CA). All the experiments were approved by and conducted in accordance with the guidelines provided by AALAC. Adult zebrafish were placed in a mating tank one day prior to embryo collection for breeding purposes. The facility was maintained at a temperature of 28 deg C along with a 14/10-hour light/dark cycle. For all the experiments, 6-7 days post fertilization (dpf) zebrafish larvae were used. Motor activity and heart rate analysis involved wild type strain (abwt), and a transgenic zebrafish strain (mnx1:GCaMP) expressing fluorescence via calcium indicator (GCaMP) in motor neurons was used for recording motor neuron activity. Breeding tanks were set with adult fish one day prior to egg collection later in the evening. Embryos were collected post-breeding and kept in the fish facility in a petri dish in fish water (conductivity: 600-750, pH: 7.2-7.8). The fish water was changed every day until the larvae hatched at 3-4dpf and kept until they turned 6 dpf. Food-restricted groups were kept food-deprived right from birth, and fed groups were given food once they turned 5 dpf, and both groups were kept separately in the facility. Larvae were taken to a separate research facility where experiments were conducted.

### Motor activity recording setup

Zebrafish Larvae (6-7dpf) were brought from the fish facility to the research facility for motor activity recording and analysis in the morning. For measuring total distances travelled by zebrafish larvae, both food-restricted and fed larvae were separated into a total of 4 groups altogether, fed control and food-restricted control (i.e., control groups) and fed with AMPH and food-restricted with AMPH (i.e., AMPH treated groups) (Fig, 1A). In this study, the terms AMPH, amphetamine, and d-amphetamine are used interchangeably.

Control groups and AMPH treated groups were initially kept in a 48-well plate (for locomotor recording) in fish water. Control larval group was not administered AMPH throughout the 20 min recording time whereas AMPH treated was recorded for first 10 mins without AMPH in fish water and then fish pipetted was out and replaced with 0.5ml of 0.7uM d-amphetamine AMPH solution (Sigma Aldrich, MO). Recording apparatus consisted of Kaiser camera stand and Basler Gencam1 GigE camera equipped with Computar lens with specifications 4.5-12mm, IR 1/2” setup in an inverted position on the camera stand (Noldus Information Technology, VA, USA). Dissection stereoscope with the in-built light source was modified by removing the eyepiece section for convenient recording, and it was used as a stage to place the well plate and petri dish (Fig. 2A). According to a study, larval zebrafish display heightened locomotor response in the darker environment (Basnet); hence, an optcast infrared filter screen 4” x 5” (43954, Edmund Optics, NJ, USA) was placed between the light source and larvae.

### Motor activity recording procedure

For locomotor activity recording, larvae placed in a 48-well plate were moved in the recording setup on the IR filter optcast that covered the light source and allowed for habituation for 5-7 mins before recording was started. Since the well plates were circular and were causing distortion from the sides of the wells, a screen of Fresnel lens with dimensions 6.7” x 6.7” (46614, Edmund Optics, NJ, USA) was placed on the plate. Fresnel lens reduced the distortion and increased the light-capturing capability of the camera by focusing maximum light into the camera lens from surrounding wells. After larval placement, recordings were commenced first in control groups for 20 mins in fish water without AMPH addition. For AMPH treated groups, larvae were similarly placed as control groups in the setup and recording was first performed for 10 mins in fish water in the absence of AMPH at 30 fps. The recording was stopped after 10 mins, and fish water was replaced entirely by AMPH solution and larvae were allowed to absorb or consume AMPH for 2-3 mins, and recording was resumed for 10 more minutes. Recordings of fed larval groups (control and AMPH treated) were carried out in the same way as it was done in food-restricted groups. Recording, tracking, and analysis of larval swimming motion was performed using Ethovision XT 15 software (Noldus Information Technology, VA, USA) which was connected to the camera in the recording setup (Fig. 2A). From the software, excel data files were generated, and data from the excel files were sorted and exported to OriginPro 2021 (OriginLab Corporation, MA) software for statistical analysis.

### Post Processing in motor activities

Recorded video was automatically converted into data for total distances travelled, and active time was by Noldus Ethovision software and downloaded as excel spreadsheets. Data from the spreadsheets were sorted, and distance data were divided by active time data to obtain velocity data points. All the data were exported to OriginPro for statistical analysis.

### Heart rate and motor neuronal activity recording setup

Similar to motor activities, food-restricted and sated larvae were divided into control groups (recording without AMPH exposure for 20 mins) and AMPH treated groups (recording without AMPH for first 10 mins and with AMPH for next 10 mins). Heart rate activity was recorded in a bright-field setup (an inverted microscope with a built-in light source for field illumination), and motor neuron activity was recorded using epifluorescence setup (fluorescence excitation and illumination setup (X-cite 120 series, Excelitas Technologies Corp., Canada), a GFP filter with ex: 488nm/em: 507nm (Chroma Technology Corp, VT)) on the same inverted microscope (Olympus Corporation, Japan). The inverted microscope was mounted with a high-speed sCMOS camera (Hamamatsu Orca Flash 4.0, Hamamatsu Photonics, Japan). A water immersion objective, 40x/0.80w LUMPlanFI (Olympus Corporation, Japan), was used to visualize and record heart and neuronal activity in the larva. The camera was connected to a live imaging visualization software (HCI Image Live, Hamamatsu Photonics, Japan) that was used to record heart and neuronal activity at 35fps with 10ms exposure time.

To record heart activity, abwt wild type zebrafish larvae were first paralyzed using 20ul of 300uM pancuronium bromide (P1918, 10MG, Millipore Sigma, WI, USA) to avoid unnecessary movements such as twitching which could interfere with the recording process. Paralysis was followed by complete immobilization of larva in a drop of 1.5% agarose gel drop and placed in a petri dish where it was covered with 300ul of fish water. The dish was kept on the microscope stage, and was larva was illuminated with an in-built light source from the bottom of the microscope. The recording in the control group was performed by covering the larva immobilized in the gel with fish water for 20 mins without administering the AMPH. AMPH treated groups were recorded in fish water for the first 10 mins without AMPH. The fish water was pipetted out and replaced with 0.3 ml of 0.7uM d-amphetamine and waited for 3-4 mins to allow the AMPH solution to penetrate the gel and absorbed by the larval skin and mouth and recording the continued again to 10 more minutes.

Motor neuronal activity was recorded in mnx1:GCaMP zebrafish larvae by paralyzing and completely immobilizing them in agarose gel (similar to heart activity). Neuronal activity was recorded only in the AMPH treated group for 40 mins. The recording was started with immobilized larva covered in fish water for 20 mins without AMPH. After the first 20 mins recording, fish water was pipetted out and covered with AMPH solution, and AMPH was allowed to penetrate the gel and reach the larva for absorption and recording was again performed for 20 mins with AMPH.

### Post-Processing of heart rate and neuronal activity

Heart activity recording was obtained from HCI live imaging in tiff file format. Tiff files were opened in ImageJ, and ROI was selected using a circular segment in the activity area. Activity data were collected from ImageJ ROI and exported to Microsoft Excel and OriginPro for the frame to time conversion and constructing time vs intensity plots. Plots were smoothened using adjacent averaging with a weighted average (window points = 4). Smoothened plots were analyzed using peak analyzer function and find peaks options were used to find baseline with asymmetric least squares smoothing count the peaks automatically (local maximum, local points = 5, height threshold = 20%).

Neuronal activity recording was obtained from HCI Image Live as tiff files and opened in ImageJ. Circular ROIs were selected from two regions, one from the neuron itself and the other from a noisy region. The data were exported to Microsoft Excel to convert frames into time and subtract neurons’ intensity values from noise values to obtain exact neuronal intensity values for sharper peaks. Data were then exported to Origin pro, where the peaks were further smoothened using adjacent averaging with weighted average and window points = 6. Baseline was obtained from post-smoothened data using asymmetric least squares smoothing. Baseline data were subtracted from smoothened peak data, and the result was divided by the baseline to obtain normalized peak data. Peaks were counted from normalized data with a threshold of 10%, local maximum with local points = 3 and peak count data were statistically analyzed.

### Statistical analysis

A normality test was conducted for all data for all activities, and a statistical test was selected accordingly. For normally distributed data, paired t-tests and two-sample t-tests were used, and for skewed distributions, Wilcoxon signed rank test and Mann-Whitney test was conducted. All tests were performed with significance (n.s. at p > 0.05, * p < 0.05, ** p < 0.01, *** p < 0.001) and represented with SEM bars.

## Notes

### Competing Interest Statement

The authors have declared no competing interest.

